# Immunization with Plant-based Vaccine Expressing *Toxoplasma gondii* SAG1 Fused to Plant HSP90 Elicits Protective Immune Response in Lambs

**DOI:** 10.1101/2024.11.15.623790

**Authors:** Lucía M. Campero, Ignacio Gual, Valeria A. Sander, Luisa F. Mendoza Morales, Victor A. Ramos Duarte, Paula M. Formigo, Emiliano Sosa, Fermín Lázaro, María Valeria Scioli, Agustín Atela, Ariel Legarralde, Federico A. Hozbor, Germán J. Cantón, Sergio O. Angel, Dadín P. Moore, Marina Clemente

## Abstract

*Toxoplasma gondii* is a protozoan parasite causing toxoplasmosis, a principal concern for public health and livestock industries. Effective vaccination strategies are crucial for controlling toxoplasmosis, particularly in the lamb, which are significant reservoirs of *T. gondii*. In addition, ovine toxoplasmosis also causes economic losses due to abortions and reproductive complications. In this study, we evaluated two immunization strategies to elucidate the immune protective potential of SAG1 fused to the plant Hsp90 adjuvant against experimental toxoplasmosis in lambs. We performed an oral administration of AtHsp81.2–SAG1HC-infiltrated fresh leaves homogenate (Plant Vaccine) and a subcutaneous administration of recombinant NbHsp90.3-SAG1HC produced in *Escherichia coli* (Recombinant Vaccine). Our results showed that only the Recombinant Vaccine significantly increased anti-rSAG1 total IgG values. In addition, only lambs immunized with the Plant Vaccine showed a significant increase in IFN-γ serum levels after the experimental infection (evaluated 8 days post-challenge). On the other hand, we also observed a statistically significant decrease in histopathological lesions (injury score) in challenged vaccinated lambs compared to challenged but not vaccinated animals (vehicle and control groups). Finally, to evaluate *T. gondii* infection, we choose the chimera rGra4-Gra7 as an acute phase protein marker. All lambs from the control and vehicle groups showed higher rates of serological reactivity than lambs from the vaccinated groups, concurrently with increased severity of lesions. These results suggest that both the plant-based and recombinant vaccines are promising candidates for controlling *T. gondii* infection in lambs, with potential benefits for enhancing public health and animal welfare.

## 1. Introduction

Toxoplasmosis is an infection caused by the protozoan *Toxoplasma gondii*, capable of infecting any warm-blooded animal. This infection may be postnatally or congenitally acquired [1]. Postnatal infection (horizontal route) occurs by ingestion of *T. gondii* oocysts present in the environment (water and soil) or by the consumption of tissue cysts present in raw or undercooked meat from infected hosts [2,3]. Congenital infection (vertical route) occurs when *T. gondii* tachyzoites cross the placentae of the dams, infecting the fetus. In humans, *T. gondii* infection is generally asymptomatic; however, in children with congenital infection or immunocompromised adults, it causes severe clinical manifestations, mostly involving the central nervous system and ocular disease [4,5].

*Toxoplasma gondii* infection in farm animals represents a risk to public health since contaminated meat consumption is one of the principal sources of human toxoplasmosis. The total economic impact of foodborne toxoplasmosis was estimated to be US$3,456 million, mainly related to healthcare costs [6]. Small ruminants (goats and sheep) and pigs are the principal sources of *T. gondii* infection for humans, also representing the principal hosts of this parasite in some countries [3]. Particularly in Argentina, the sheep industry is of high socio-economic importance, and according to Instituto Nacional de Tecnología Agropecuria (INTA), Argentina has an estimated number of 15 million sheep destined mainly for meat and wool production. However, sheep dairy farming is also increasing. *Toxoplasma gondii* infection rate in sheep flocks in Argentina is near 10% [7], while in other regions it ranges between 1.4% and 98% [8]. Besides *T. gondii* foodborne zoonotic infections, they also significantly impact animal production. *Toxoplasma gondii* infection one of the most common causes of abortions in sheep and goats worldwide [9]. Ovine toxoplasmosis is also responsible for 1-2% of neonatal losses per year in the United Kingdom [10]. Additionally, sheep infected with *T. gondii* during pregnancy can vertically transmit the parasite to the offspring, contributing to the spread of the disease [8].

Given the high prevalence of ovine toxoplasmosis worldwide and its risk to public health, *T. gondii* infection control is of relevance. Vaccination would be the most successful and economical route for limiting its propagation [11,12]. Although live vaccines have shown to be efficient, they require expensive production systems, cold chain transport, and appropriate handling. On the other hand, recombinant vaccines overcome all these problems, but in general, they do not generate an immune response as robust as live vaccines. Several reports have tested recombinant vaccines conferring immune response in sheep; however, in almost all cases, they did not test their efficiencies after *T. gondii* challenge [13]. As an exception, Supply et al. [14] developed a BCG strain that produces and secretes the *T. gondii* dense granule protein GRA1 administrated subcutaneously in sheep. After an oral challenge with *T. gondii* oocysts, they evaluated the vaccine efficiency by measuring rectal temperature and observed partial protection in six animals.

Our laboratory has developed and evaluated different plant-based vaccine systems against *T. gondii* using the mouse model [15–18]. We observed that the evaluated vaccines triggered a Th1 or Th1/Th2 immune response and conferred partial protection against *T. gondii* cyst formation. In addition, we demonstrated that two isoforms of the family of plant heat shock proteins 90-kDa (pHsp90) can modulate and also enhance the immune response when they are used as carriers/adjuvants [18–20]. In our experience, the fusion of the antigen of interest to pHsp90s also improves the antigen expression [20]. The expression of antigens in plants would, in turn, generate a safe immunogen, and the use of plant extracts expressing the antigen of interest protects it from digestive degradation [20,21]. In particular, we generated a fusion protein including a B- and T-cell antigenic epitope-containing surface protein SAG1 (SAG1HC) fused to the Hsp90 chaperone of *Arabidopsis thaliana* (AtHsp81.2-SAG1HC). The mice immunizations with plant extracts from AtHsp81.2–SAG1HC-infiltrated fresh *Nicotiana benthamiana* leaves showed high efficiency in eliciting a protective immune response, reducing about 20 times the number of brain *T. gondii* cysts compared to the non-immunized control group after *T. gondii* challenge [20]. The AtHsp81.2–SAG1HC-infiltrated fresh leaves showed protective properties against *T. gondii*, facilitating oral immunization. In addition, it also presents some adjuvant components that would stimulate the immune response [17]. Additionally, the AtHsp81.2 and other members of the plant Hsp90 (pHsp90) family have shown adjuvant capabilities on their own, which makes pHsp90s attractive candidates as vaccine adjuvants against intracellular pathogens [16,22].

To know the immunoprotective effect of SAG1 fused to the pHsp90 adjuvant against acquired toxoplasmosis in lambs, we evaluated two immunization strategies: i. Oral administration of AtHsp81.2–SAG1HC-infiltrated fresh leaves homogenate (Plant Vaccine) and ii. Subcutaneous administration of recombinant *N. benthamiana* Hsp90.3 fused to SAG1HC (NbHsp90.3-SAG1HC) purified from a pRSETA-NbHsp90.3-SAG1HC plasmid expressed in *Escherichia coli* (Recombinant Vaccine). Our results showed that immunized lambs from the Recombinant Vaccine group showed an increase in the anti-rSAG1 humoral response following the third immunization. In addition, after subcutaneous challenge with *T. gondii* tachyzoites from the RH strain, lambs immunized with Plant Vaccine showed significantly higher IFN-γ serum levels when compared to controls. On the other hand, immunized lambs from the Recombinant Vaccine and Plant Vaccine groups showed diminished lesion scores and reduced brain cyst loading compared to control infected lambs. We discussed the value of these vaccine candidates against *T. gondii* in lambs of each immunization system.

## 2. Materials and Methods

### 2.1. Parasites

The RHΔhxgprt strain [23] was grown and kept under standard tachyzoite culture conditions. Monolayers of hTERT line fibroblasts (ATCC® CRL-4001, USA) were infected with tachyzoites, incubated with Dulbecco’s modified Eagle’s medium (DMEM, Invitrogen) supplemented with 1% fetal bovine serum (FBS, Internegocios S.A., Argentina) and penicillin (10,000 IU/ml)-streptomycin (10 mg/ml) solution (Gibco, Argentina) at 37°C and 5% CO_2_.

### 2.2. Recombinant SAG1HC fused to plant HSP90

A fragment (from residue 221 to residue 319 of the protein sequence) of the *T. gondii* surface protein, also known as P30 [24,25] or SRS29B (TGME49_233460), denominated SAG1HC [18] was used.

For the immunization trial, two versions of SAG1HC were used, in both cases fused to plant Hsp90. In a first version, SAG1HC was fused to NbHsp90.3 and the addition of a tag encoding 6 histidine residues was also performed to obtain the product 6His-NbHsp90.3-SAG1HC, from now on NbHsp90.3-SAG1HC. This recombinant antigen NbHsp90.3-SAG1HC was expressed and purified from *E. coli* culture as previously described by Sánchez-López et al. [18]. The purified protein was electrophoresed on SDS-PAGE and stained with Coomassie blue (Fig. S1). The protein was quantified by the Bradford method. The purified product was used as immunogen (Recombinant Vaccine).

In the second version, SAG1HC was fused to AtHsp82.1 to obtain the expression product 6His-AtHsp81.2-SAG1HC, and from now on AtHsp81.2-SAG1HC. The protein was transiently expressed in *N. benthamiana* leaves using *Agrobacterium thumefaciens* infiltration as described by Sánchez-López et al. [20] and quantified by Western blot using the Gel-Pro analyzer software (Media Cybernetics, Rockville, MD, United States) (Fig. S1) as previously described [20]. In this case, the plant extract from AtHsp81.2–SAG1HC-infiltrated fresh leaves were used as an immunogen (Plant Vaccine).

### 2.3. Immunization and Challenge

Fifteen seronegative three-month-old Texel lambs were selected for the experiment. The lambs belong to a flock at Instituto Nacional de Tecnología Agropecuaria (INTA), Balcarce. They were kept in closed pens and received commercial alfalfa pellets and *ad libitum* water. They were bled before the beginning of the immunization protocol and their seronegative status to *T. gondii* and *Neospora caninum* was confirmed by indirect fluorescence antibody test (IFAT; cut-off titer of ≥1:50) [26]. The animals were in good body condition and their health was monitored throughout the experiment. Vaccination and sampling schedule are summarized in Fig. 1A. Briefly, lambs were randomly allocated into 4 groups: Control (PBS-immunized lambs, *n* = 4); Vehicle (lambs orally immunized with non-infiltrated fresh leaves, *n* = 4); Recombinant Vaccine (lambs subcutaneously immunized with recombinant NbHsp90.3-SAG1HC produced in bacteria, *n* = 3), and Plant Vaccine (lambs orally immunized with AtHsp81.2–SAG1HC-infiltrated fresh leaves, *n* = 4).

**Fig 1.**
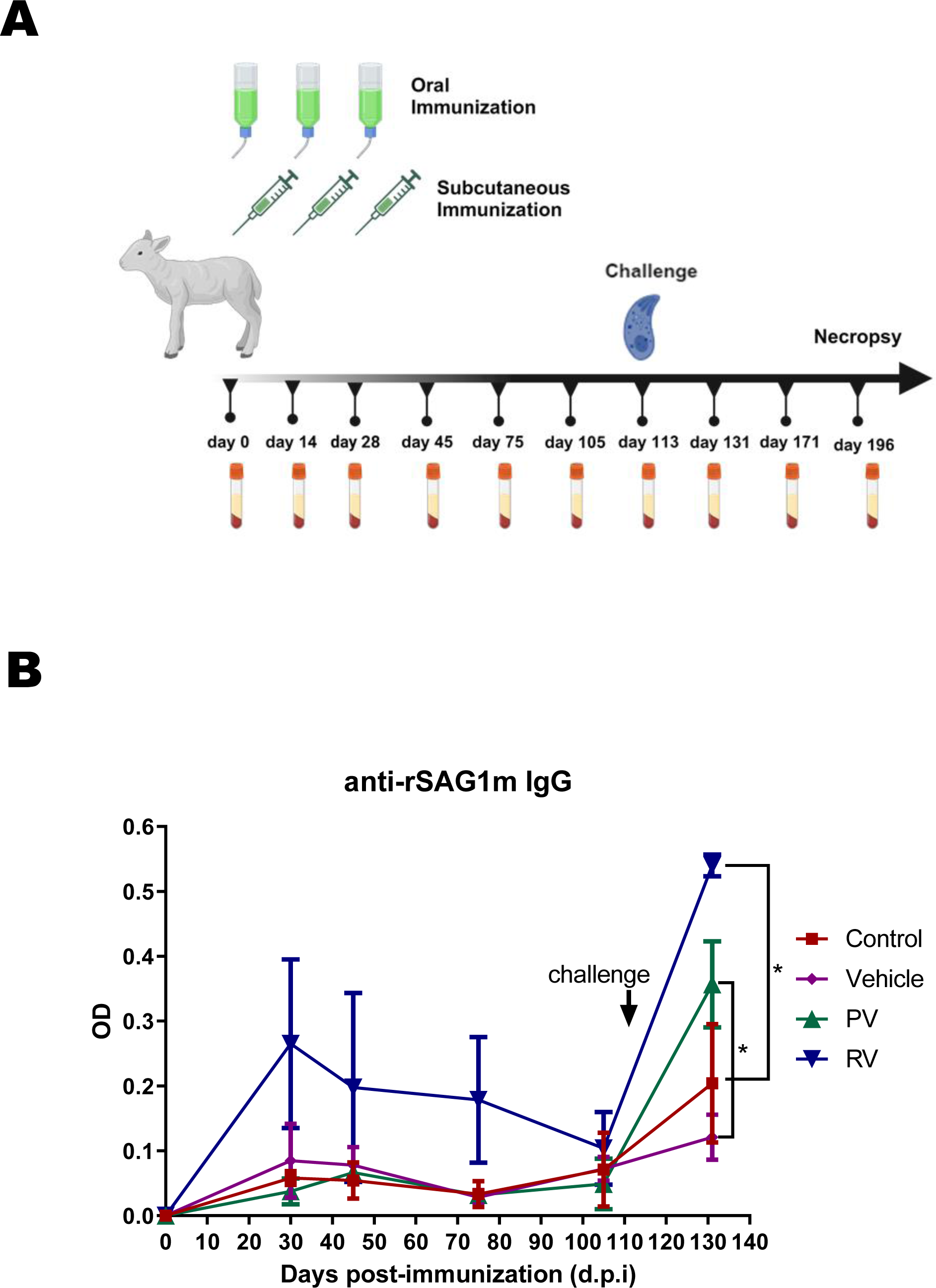
Immunization of lambs with the SAG1HC antigen fused to plant HSP90. A. Immunization plan and challenge protocol. Recombinant Vaccine: subcutaneously immunized lambs with the recombinant NbHsp90-SAG1HC purified from *E. coli*; Plant Vaccine: orally immunized lambs with AtHsp81.2-SAG1HC-infiltrated fresh leaves; Control group: orally immunized lambs with PBS buffer; Vehicle group: orally immunized lambs with non-infiltrated fresh leaves. All Lambs received three doses separated by 14 days. Blood samples were drawn to monitor the kinetics of specific antibody production. All lambs were subcutaneously challenged with 5 x 10^6^ tachyzoites from the *T. gondii* RH strain. On day 196, the lambs were euthanized, and blood and tissue samples were obtained. B. The antibody kinetics monitoring in sera from Control, Vehicle, Recombinant Vaccine, and Plant Vaccine groups against rSAG1 purified from *E. coli*. Statistical analyses were performed using two-way ANOVA followed by Tukey’s multiple comparisons test. *p < 0.05. The graph is representative of two independent experiments with similar results.

Vaccine formulations were as follows: Vehicle, 38 g of tobacco leaf extract homogenized in 200 ml of PBS per dose; Recombinant Vaccine, 0.3 mg of rNbHSp90.3-SAG1HC in 3 ml of PBS per dose; and Plant Vaccine, 38 g of infiltrated fresh leaves containing 0.5 mg of pHsp81.2-SAG1HC homogenized in 200 ml of PBS per dose. None of the vaccine formulations evaluated here included any additional external adjuvant. Local inflammatory reactions at the injection site were evaluated daily for one week after inoculation, visually and by palpation. Blood samples from each animal (10 ml) were collected from the jugular vein for serological assays at days 0, 14, 28, 45, 75, 105, 113, 121, 131, 171, and 196 days post-immunization (d.p.i.). Serum was obtained by centrifugation (1,000 g, 15 m, RT).

Lambs from Control, Vehicle, Recombinant Vaccine and Plant Vaccine groups were subcutaneously challenged with 5 x 10^6^ *T. gondii* tachyzoites from the RH strain at 113 d.p.i. [27]. After field inoculation, an aliquot with the parasite dose challenge was used to infect hTERT cells to evaluate viability in field conditions.

All animals used in this study were handled in strict accordance with good animal practice and the conditions defined by the Animal Ethics Committee at INTA Balcarce (CICUAE 244/2022).

### 2.4. *Escherichia coli*-Recombinant antigens, *Toxoplasma* lysate antigens (TLA) and Serology

The rSAG1 protein was purified as described [18]. The rGra4-Gra7 chimera was obtained from the gene sequences that correspond to the C-terminal region of rGra4_163-345_ and the mature form of the native protein of rGra7_27- 236_, which were fused with a linker (Gly-Ser-Gly-Ser-Gly) region between Gra4 and Gra7 fragments [28]. The rGra4-Gra7 fusion protein also encodes 6 histidine residues at the N-terminus. The complete sequence was synthesized, and the product was cloned between the NdeI/BamHI sites of the expression vector pET-14b (GenScript, Piscataway, NJ, USA). The rSAG1 protein from *E. coli* was obtained as previously described by [18]. Tachyzoite lysate antigen (TLA) was prepared from purified RH tachyzoites grown in HFF culture. After purification by a 3 µm polycarbonate filter, tachyzoites were sonicated as described by Sánchez-López et al. (2019). Bradford protein assay was used to quantify the TLA and then, it was stored at -80 °C.

All recombinant proteins were purified from *E. coli* cultures induced with 0.5 mM IPTG overnight at 37°C with shaking. Purification was carried out by Nickel affinity chromatography using a Ni-NTA resin (Ni2+-nitrilotriacetic acid resin, Qiagen, Valencia, CA, USA) following the manufacturer’s recommendations under denaturing conditions.

Polystyrene 96-well microtitre plates (Nalge Nunc International) were coated overnight with 5 μg/ml of different recombinant proteins diluted in 50 mM carbonate buffer (pH 9.6), blocked for 1 h at 37°C with blocking solution (5% milk in TBS-Tween: 100 mM Tris–HCl, pH 7.5, 0.9% NaCl, 0.1% Tween 20), and washed three times with TBS–Tween. After blocking, 100 μl of each serum diluted 1:100 in the blocking solution were added to each well in duplicates and were incubated for 1 h at 37°C. After that, plates were washed three times. One hundred μl of a 1/5,000 dilution of horseradish peroxidase-conjugated goat anti-sheep IgG (Jackson ImmunoResearch) diluted in blocking solution was added to each well, and incubated for 60 min at 37°C. After three washes, immune complexes were revealed with tetramethylbenzidine chromogen (TMB, One-Step; Invitrogen, Carlsbad, CA, USA) and optical density (OD) was read at 630 nm with an automatic ELISA reader (Synergy H1, Bio-Tek, VT, USA). Ten serum samples of negative sheep determined by indirect immunofluorescent antibody test (IFAT) and TLA-based IgG-ELISA were used as the negative control. A serum sample was considered positive if the OD value was higher than the cut-off value (mean OD value for negative control plus 3 standard deviations). As a positive control, a pool of sera from *T. gondii* seropositive sheep characterized by IFAT was used. Each serum sample was examined twice in duplicate.

For titer analysis, sera from animals corresponding to each group were pooled as well as sera from 10 seronegative animals for each time point. Serial dilutions from 1:100, were analyzed by IgG-ELISA. The highest dilution that still gave a positive result was taken as the titer value.

### 2.5. Cellular immune response: cytokines and chemokine levels

The levels of IFN-γ, IL-12 and IL-4 cytokines, as well as the chemokine IP-10 (interferon-induced protein of 10-kDa) were determined using ovine IFN-γ, IL-12, IL-4 and IP-10 ELISA Kits (Abcam Inc, MA, USA), according to the manufacturer’s recommendations. Serum samples from each animal corresponding to the different groups were obtained before (0 d.p.i.) and after (121 d.p.i., 8 days post-challenge) the experimental infection with *T. gondii*. A 1:2 dilution of each sample was used in the tests. Cytokines and chemokine concentrations were estimated by interpolation from standard curves based on recombinant cytokines/chemokine included in the corresponding ELISA kit. Concentration values were only considered when inside the detection limits indicated by the manufacturer.

### 2.6. Euthanasia, necropsy and histopathological analysis

Lams from Control, Vehicle, Recombinant Vaccine and Plant Vaccine groups were euthanized 77 days post-challenge, following the recommendations of CICUAE 244/2022. Briefly, animals were stunned with a captive bolt then immediately jugular bleed out. Post-mortem examination was carried out immediately after confirming death. Tissues were fixed in 10% buffered formalin solution for 5 days and later processed by standard histopathological methodologies for hematoxylin and eosin staining. Tissue slides were randomized, blind-coded and examined under optical microscopy conditions using various magnifications (10×, 20× and 40×). The scores were obtained by the two blind independent observers and an average was calculated. A scoring range of 0 to 3 was used where 0 indicated the absence of inflammatory infiltrate and necrosis; 1 denoted focal infiltration of inflammatory cells; 2 represented multiple foci of moderate inflammatory infiltrate; and 3 indicated diffuse inflammatory infiltration accompanied by edema and necrosis.

## 2.7. Statistical analysis

Statistical analyses were carried out using GraphPad Prism 5.0 Software (GraphPad, CA, USA). Two-way analysis of variance (ANOVA) or Two-way analysis of variance with repeated measures (RM), followed by Tukey’s multiple comparison tests were used to compare variables from experimental groups, p < 0.05 was considered as statistically significant.

## 3. Results

### 3.1. Immunization

Previously, we have studied the adjuvant capacity of pHSP90s to stimulate the immune response of different antigens of interest [22]. In addition, we have demonstrated that AtHsp81.2-SAG1HC as an oral Plant Vaccine has immunoprotective value against toxoplasmic infection in mice [20], while the subcutaneous administration of recombinant NbHsp90.3-SAG1HC was immunoprotective in experimentally-infected mice [18]. To analyze the protective value of both platforms in lambs, here we design a similar strategy: one with AtHsp81.2–SAG1HC-infiltrated *N. benthamiana* fresh leaves (Plant Vaccine), and the other with the recombinant NbHsp90.3-SAG1HC protein purified from *E. coli* (Recombinant Vaccine) (Fig. 1A).

Before the challenge (during the monitoring of the lambs) a significant increase in the anti-rSAG1 IgG OD values was shown only in the lambs of Recombinant Vaccine group (Fig. 1B). Post-challenge, all lambs from each group showed an increase in the humoral immune response against rSAG1 (Fig. 1B). To note, lambs from Plant Vaccine and Recombinant Vaccine groups showed OD values significantly higher than the corresponding controls (Vehicle and Control groups, respectively).

### 3.2. Humoral immune response analysis

To check if any animal had a pre-challenge *T. gondii* infection, the sera from day 105 d.p.i. of all groups were analyzed by anti-rGra4-Gra7 IgG-ELISA. The IgG-ELISA analysis demonstrated that one of the lambs belonging to the Control group elicited a humoral response well above the cutoff point (Fig. S2). Therefore, we decided to discard this lamb for subsequent analysis.

The experimental infection resulted in IgG levels above the cutoff levels in all lambs from each group, as indicated by the analysis of specific antibodies against total lysate antigens (TLA) from *T. gondii* tachyzoites (Fig 2). However, the results showed differences between groups regarding the intensity or the profile of the immune response. After the challenge, lambs from the Vehicle group showed greater reactivity than the remaining groups (Control, Recombinant Vaccine and Plant Vaccine).

**Fig. 2.**
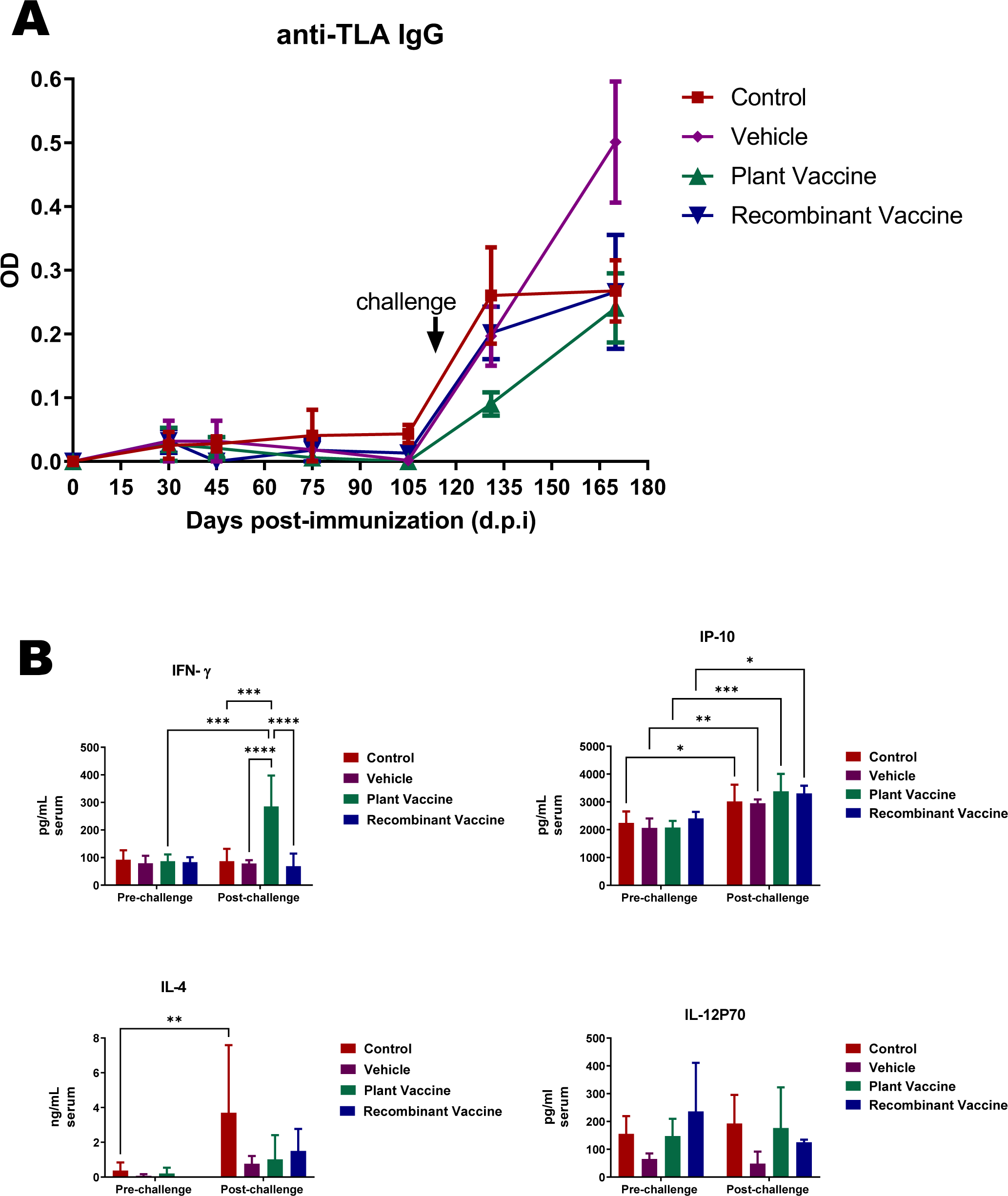
Immune response of lambs. A. The serology against *T. gondii* of the immunized and challenged lambs was analyzed by the IgG-ELISA technique using TLA as an antigen. Challenge (113 d.p.i.) is indicated. The graph is representative of two independent experiments with similar results. B. Cytokines were analyzed in serum samples obtained from animals on day 0 (pre-challenge) and day 121 (post-challenge) of the experiment, with the latter representing day 8 post-infection (acute infection). Statistical analyses were performed using Two-way ANOVA with repeated measures (RM), followed by Tukey’s multiple comparisons test. *p < 0.05, **p < 0.01, ***p < 0.001 and ****p<0.0001. The graph is representative of two independent experiments with similar results.

### 3.3. Cellular immune response analysis: cytokine and chemokine levels

The success of vaccination against *T. gondii* involves triggering a cellular immune response, with IFN-γ playing a key role [29]. It is known that IL-12 and the chemokine IP-10 (interferon-induced protein of 10 kDa) enhance IFN-γ secretion [30,31], whereas IL-4 inhibits its production [32]. Taking these data into account, we decided to analyze serum cytokine levels at day 0 d.p.i. (pre-challenge) and 121 d.p.i. (8 days post-challenge, acute infection) [33]. As can be observed in Fig 2B, no differences in IFN-γ levels between groups were found at 0 d.p.i.; however, lambs from the Plant Vaccine group showed significantly higher levels of IFN-γ during acute infection (8 days post-challenge, 121 d.p.i.) when compared with lambs of other groups (Control, Vehicle and Recombinant Vaccine) and also with levels of IFN-γ secreted by the same animals from all groups before experimental infection. Additionally, there was a significant increase in IP-10 levels in all groups 8 days post-challenge, being the lambs of Plant Vaccine group the one that showed the most significant increment. No differences were found in IL-12p40 levels between groups at 0 d.p.i. or at 121 d.p.i., nor any increase within the same groups after experimental infection. Finally, no differences were found in IL-4 levels between groups at 0 or 121 d.p.i., neither within the same group between 0 and 121 d.p.i., with the exclusion of Control group, which showed a significant increment on 121 d.p.i. when compared with day 0 d.p.i. These results would be in agreement with the possibility of an enhancement of the Th1-like cellular immune response in the lambs of Plant Vaccine group after experimental infection (8 days post-challenge).

### 3.4. Analysis of the protective value of vaccines

*Toxoplasma gondii* can rapidly disseminate throughout all the tissues, where it replicates before passing into its dormant state of encysted bradyzoite. This process generates visible tissue lesions and inflammatory processes. Given the lack of visible clinical signs, we analyzed the protective value of different vaccines against *T. gondii* infection by evaluating the severity of injury produced mainly in brain samples [34]. Taking this data into account, we analyzed the histopathological lesions from different brain regions of the lambs euthanized 77 days post-challenge. Table S1 shows the results obtained. Figure S3 shows photomicrographs of brain tissues showing different histopathological lesions and the corresponding assigned score.

The injury score found in lambs from the Recombinant Vaccine or the Plant Vaccine groups showed a significant reduction when compared with lambs from Vehicle or Control groups, suggesting that both formulations partially protected lambs from the *T. gondii* challenge. Additionally, all analyzed brain regions showed lesions and inflammation that could be attributed to the infection (Fig. 3B).

**Fig. 3.**
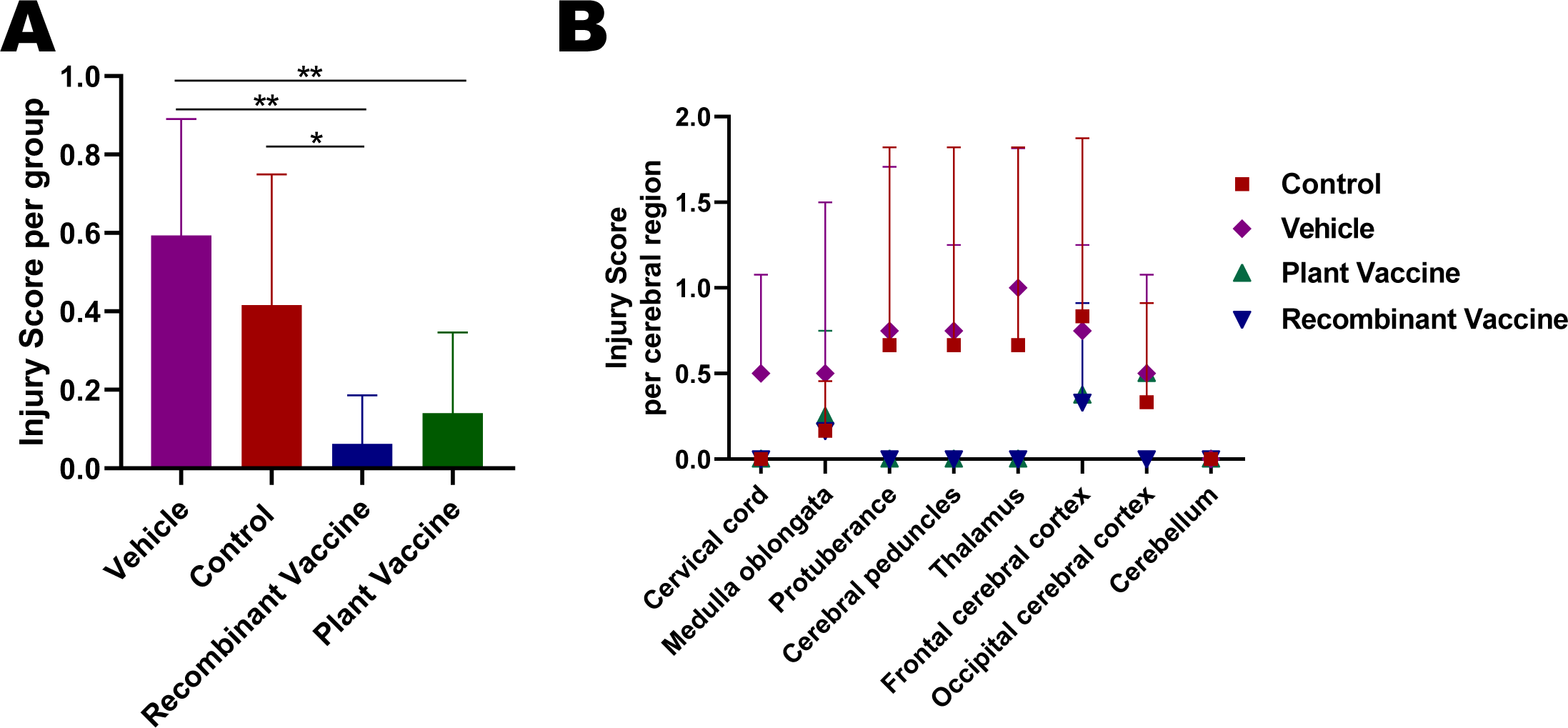
Histopathological analysis in tissue samples from *Toxoplasma gondii* challenged lambs. **A.** Analysis of the brain injury scores per group. The lesion and inflammation scores were ranked from 0.5 (mild), 1 (moderate), to 2 (severe) (see Table S1). **B.** Analysis of the lesion scores by brain area. The lesion score values depicted for each experimental group are expressed as the mean value + S.D. The statistical analysis was carried out for Tukey’s multiple comparisons test.

### 3.5. Cyst and PCR detection

A significant aspect of an anti-*T. gondii* vaccine in lambs is to reduce encystment to control foodborne-associated infections. In our hands, the *T. gondii* cysts visualization from brain samples by optical microscopy was arduous, even after purification by Percoll gradients [27]. Furthermore, the presence of *T. gondii* cysts was analyzed in lamb brains during histopathological analysis. We visualized brain *T. gondii* cysts in three lambs from the Vehicle group and one lamb from the Plant Vaccine group (Table S1). We also analyzed the presence of *T. gondii* genomic DNA (gDNA) in brain tissue by PCR as described [35], but only one sample of gDNA from *T. gondii* was detected (lamb 162 belongs to the Plant Vaccine group; data not shown).

### 3.6. Serology as marker of T. gondii infections

To analyze the acute phase outcome, we chose the rGra4-Gra7 chimera as an acute phase marker [28]. Since the sera from different animals showed different reactivity values against rGra4-Gra7, we analyzed each one of them separately. This analysis showed that the reactivity profiles were similar among groups. All lambs, except one from the control group (152), showed the anti-rGra4-Gra7 response to rise on day 131 above the cutoff, declining by day 170 (Fig. 4). Remarkably, the OD values at day 131 of the control and vehicle groups were mostly higher than those observed in the vaccinated groups (Fig. 4). Day 131 of the experiment corresponds to day 18 post-challenge, coinciding with the acute stage of *T. gondii* infection. To analyze this stage in more detail, pools of the sera from each group were made and their titers were tested (Fig. 4). As can be observed, the Control (1:6400) and Vehicle (1:6400) groups present two to three higher titers than the Recombinant Vaccine (1:800) and Plant Vaccine (1:1600) groups, respectively.

**Fig. 4.**
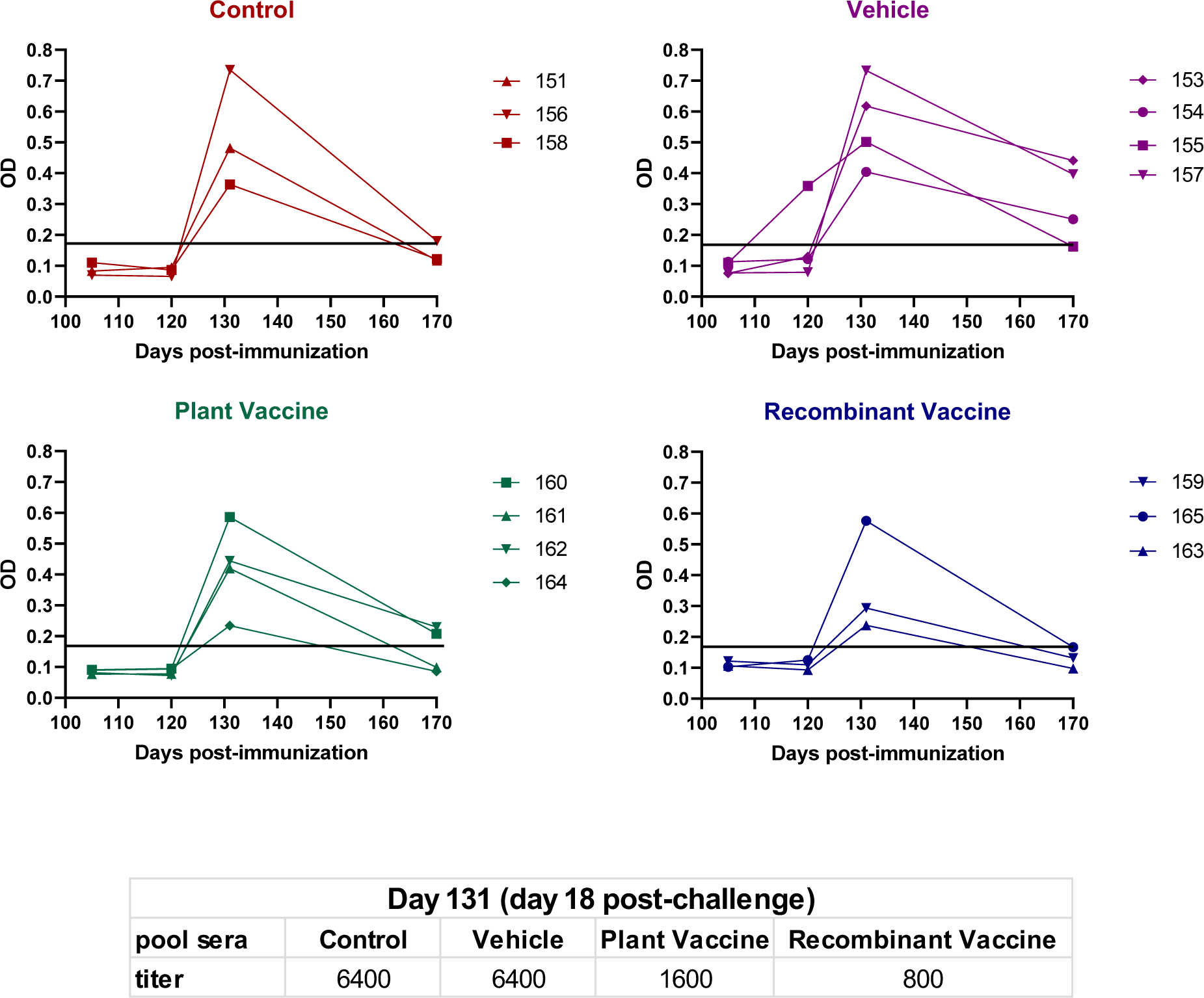
Acute infection monitoring of Toxoplasma gondii challenged lambs. The serology analysis against T. gondii of the immunized and challenged lambs by the IgG-ELISA technique using rGra4-Gra7 as an antigen. Sera were diluted at 1:100. The horizontal line indicated the cutoff (the OD of negative control serum sample + 3 S.D.). It is representative of two independent experiments with similar results. Titer of sera samples obtained at day 131 was also analyzed. The sera from each group were pooled in equal parts. Similarly, a pool of 10 negative sera was generated. A serial dilution curve was made. The titer value was obtained as the lowest dilution at which a reaction between the antibody and rGra4-Gra7 occurred.

## 4. Discussion

Experimental *T. gondii* infection in mice offers a simple model to test antigens and adjuvants in immunization plans. However, the success observed in the murine model needs to be validated in animals of interest or humans. In this work, we analyze the use of the *T. gondii* antigen SAG1HC fused to the adjuvant protein pHsp90 to immunize lambs: NbHsp90.3-SAG1HC expressed and purified from *E. coli* (Recombinant Vaccine); and a plant-based vaccine: AtHsp81.2-SAG1HC (Plant Vaccine). Both formulations showed promising results in mice [18,20]. Here, our results in lambs showed that subcutaneous immunization with NbHsp90.3-SAG1HC generated a detectable humoral response against SAG1. Post-challenge, SAG1 detection was observed in all groups, with higher O.D. in lambs from Plant Vaccine and Recombinant Vaccine groups, which suggests the enhancement of a memory B-cell response produced by immunization, as previously observed in the murine model [16,20].

It is generally accepted that control of *T. gondii* infection mainly relies on the secretion of IFN-γ, which is involved in innate and adaptive immunity [29], being considered a molecular marker of toxoplasmic infection [36]. Previous works have shown high levels of serum IFN-γ during acute experimental infection of sheep [33]. In the current study, only vaccinated lambs from the Plant Vaccine group showed a significant increase in IFN-γ serum levels after experimental challenge with *T. gondii*., thus these results could be considered contradictory. However, IFN-γ kinetics during the immune response is complex and involves the timely secretion of several mediators [10,31]. Briefly, during early infection, IL-12 is secreted mainly by activated macrophages, dendritic cells and neutrophils, and stimulates the proliferation of natural killer (NK) cells which in turn, secrete IFN-γ [33]. The early release of IFN-γ triggers the secretion of IFN-γ inducible cytokines and chemokines, including IP-10 [31]. In sheep, Buxton et al. [37] have demonstrated that the primary infection with *T. gondii* tachyzoites enhanced the secretion of IFN-γ from 2 days post-infection, reaching titer peaks 1 day later, declining as early as 6 days after infection. In our experiment, serum IFN-γ levels were determined 8 days post-infection, thus its levels (related to the innate immune response) may have declined by that time. A similar interpretation could apply to IL-12 levels, which showed no differences between pre-and post-challenge serum levels. This suggestion is supported by data on IP-10 serum levels, a cytokine secreted in response to IFN-γ [31], which were increased in every experimental group. Curiously, as previously stated, only the lambs of Plant Vaccine group showed increased levels of IFN-γ 8 days post-infection, concurrently with the most significant increase in IP-10. Noteworthy, IP-10 stimulates the recruitment of antigen-specific T cells (Th1 and TCD8+) that release IFN-γ in the sites of infection and amplify the Th1 response (Khan et al., 2000). The increment of IFN-γ 8 days post-infection in the lambs of lPlant Vaccine group suggests that a T cell-independent response became replaced by an antigen-specific T lymphocyte response around 8 d.p.i., which is in agreement with IFN-γ kinetics data from previous research in *T. gondii* experimentally infected sheep [38]. This could be related to previous vaccination with the AtHsp81.2–SAG1HC-infiltrated fresh leaves and the generation of a memory immune response. It could be in concordance with the presence of other natural T-cell-associated adjuvants present in plant extracts [17].

The results of the experimental infection with *T. gondii* show that both vaccine formulations (Recombinant Vaccine and Plant Vaccine groups) successfully generated memory immune responses in the lambs. Importantly, lambs immunized with either Recombinant Vaccine or Plant Vaccine showed protection against the parasite after experimental infection, as evidenced by a reduction in histopathological lesions. This suggests that the vaccination systems studied here can provide a partial immunoprotective response against toxoplasmic infection in sheep.

One of the objectives of our studies was to verify that plant Hsp90s can be effective adjuvants in other infection models, different from the mouse. Although AtHsp81.2 and NbHsp90.3 present high similarity at the sequence level, their immunostimulatory properties could differ [39]. Here, we observed that immunization with NbHsp90.3-SAG1HC purified from *E. coli*, without any additional adjuvant, induced a humoral immune response against SAG1 in lambs. Likewise, this formulation could be adequate to control the dissemination of *T. gondii,* analyzed as the significant decrease of inflammatory processes and tissue lesions in the brain generated during the infection. In the case of AtHsp81.2-SAG1HC, we did not observe the elicitation of a humoral response during the immunization schedule. In this case, the AtHsp81.2-SAG1HC protein was inoculated orally and present in plant extracts. However, AtHsp81.2-SAG1HC reduced the lesions and inflammatory processes after the challenge. This protection against infection correlated with high levels of IFN-γ in serum, suggesting the generation of T-cell memory immune response, as previously discussed. The differences observed in the immune response generated by the Plant vaccine and the Recombinant vaccine should not be attributed solely to the different HSP90 isoforms. It is also important to consider the impact of the various routes of administration used in this research. Although further studies are needed to elucidate the immunogenic mechanisms of both formulations, our results indicated that plant Hsp90s could be alternative adjuvants for use in the veterinary industry.

The attenuated S48 strain was widely used as a vaccine, including sheep, with high success [40,41]. However, it requires high production, transportation, and handling costs. Recombinant proteins produced in bacteria or plants are easy to scale production, more manageable, and cheaper than vaccines based on attenuated *T. gondii*. In addition, plant-based vaccines have other advantages, being safer and more efficient when administered orally [17,42]. However, obtaining an immune response such as that conferred by live *T. gondii* is a challenge. One of the most relevant difficulties is the analysis of its effectiveness. Until now, few studies have explored the response of *T. gondii* vaccines in sheep after challenge [13]. Likewise, these studies use different outcome measures to evaluate their protective efficacy against experimental infection: rate of infection in offspring, rectal temperature, and presence of cysts [12,14,27]. Based on these data, it is difficult to determine the immunoprotective efficacy of our formulations compared to other vaccines reported. Although our results are promising in the future, it is necessary to generate a study model of experimental infection in lambs to determine the efficiency of each antigen or immunization formulation.

Interestingly, the serological data are consistent with the histopathological data in the animals. In that sense, the chimera rGra4-Gra7 proved to be a feasible candidate to monitor the impact of the experimental infection during its acute stage. This chimera antigen is easy to produce and is different from the immunogen used. Therefore, it avoids cross-reactions of the vaccinated animals. Likewise, by representing two specific *T. gondii* antigens, it avoids cross-reactions due to infections of lambs with other coccidia. Interestingly, the levels of reactivity against rGra4-Gra7 were different in the Control and Vehicle groups compared to the vaccinated groups. These data could be inferring that the vaccinated groups produce a lower response against rGra4-Gra7 because the infection would be more controlled. Although more experiments are needed to confirm this trend, in this way, the reactivity against rGra4-Gra7 could be a marker of the progress of the infection compared to the control groups.

We chose the SAG1 antigen since it has been shown to be a vaccine candidate, and is also our immunization model to test the adjuvant value of plant HSP90 [43–45]. However, there are other antigens that could also be tested [46]. An important aspect is that the HSP90 protein not only functions as an adjuvant, but also as a carrier [22,47,48]. This improves protein expression, even those of low molecular weight polypeptides [20]. In the future, small *T. gondii* multi-epitope polypeptides could be expressed thus generating a broader cellular and humoral response against toxoplasmosis in comparison with the immunization of a single antigen [49–51].

## 5. Conclusion

Here, we showed that two vaccine formulations based on the SAG1 antigen fused to the plant Hsp90s used as novel adjuvants induced a humoral and/or cellular immune responses and partially reduced tissue damage cause by *T. gondii* challenge. Notably, this is the first study to assess the efficacy of a plant-based vaccine administered via oral immunization. Our findings underscore the feasibility of combining different vaccine production systems and administration routes, incorporating innovative carriers and adjuvants, such as plant Hsp90s, to develop effective vaccines for sheep.

## Author contributions

M.C. DPM and S.O.A. Writing – review & editing, Writing – original draft, Supervision, Investigation, Formal analysis, Conceptualization. L.C., I.G. and V.A.S. Methodology, Investigation, Formal analysis, Writing – review & editing. L.F.M.M, V.A.R.D., P.F., A.T., A.L., E.S., F.L., M.V.S., Methodology. G.J.C. F.A.H., Supervision, Methodology.

## Funding

The work was supported by ANPCyT (PICT 2020-639 and PICT cat. I 56), CONICET (PIP 2021-1168), MINCyT (PITEs 50), INTA (PD.I116/P02) and UNMdP (AGR717/24).

## Institutional Review Board Statement

All lambs were maintained under monitoring of natural diseases and infections and handled according to the approved institutional animal care and use committee protocols of the Animal Ethics Committee at INTA Balcarce (CICUAE 244/2022).

## Informed Consent Statement

Not applicable.

## Data Availability Statement

All data upon which conclusions are drawn are included in the manuscript.

## Acknowledgements

L.M. Campero, D.P. Moore, S.O. Angel, M. Clemente, and V.A. Sander are members of Consejo Nacional de Investigaciones Científicas y Técnicas (CONICET). S.O. Angel, M. Clemente and V.A. Sander are also Professors at Universidad Nacional General San Martin (UNSAM) and D.P. Moore is also Professor at Universidad Nacional de Mar del Plata. V.A. Gual is post-doctoral fellow of CONICET and Professor at Universidad Nacional de Mar del Plata. V.A. Ramos-Duarte and L.F. Mendoza-Morales are doctoral fellows of CONICET, and L.F. Mendoza-Morales is also a teaching assistant at UNSAM. A. Legarralde is a technical staff of CONICET. P. Formigo and A. Atela are students of UNSAM. G.J. Cantón is a researcher at the National Institute of Agricultural Technology (INTA Balcarce). We thank Sergio Dinino and Yolanda Marron for their collaboration in the necropsy. Also, we thank Agustina Ganuza, Patricia Uchiya and Jose Luis Burgos for technical assistance.

## Conflicts of Interest

The authors declare no competing interests.

## Supplementary files

**Fig. S1**. Expression of pHSP90-SAG1HC proteins. **A.** NbHsp90-SAG1HC obtained from E. coli (Recombinant vaccine) by Ni-NTA affinity and analyzed by SDS-PAGE stained with Coomassie blue. The expected band of 110-kDa is observed along with other protein degradation products. **B.** Western blot analysis and quantification with rSAG1 purified from recombinant E. coli of the AtHsp81.2-SAG1HC expression present in plant leaf extracts.

**Fig. S3.** Histopathological analysis of brains of *Toxoplasma gondii* challenged lambs. Brain tissue samples were analyzed for the presence of lesions, inflammatory processes, and the presence of *T. gondii* cysts (see Table S1). The different lesions and inflammatory processes were used to establish a score from 0 to 3. An example of the employed score is shown in the panels.

**Table S1.** Description of the histopathological lesions and presence of *T. gondii* cysts observed in the different areas of the brain per lamb from the different experimental groups.

## Notes

### Competing Interest Statement

The authors have declared no competing interest.

