## Supplementary figures and images for "Immunization with Plant-based Vaccine Expressing *Toxoplasma gondii* SAG1 Fused to Plant HSP90 Elicits Protective Immune Response in Lambs"

### Supplemental Figure 1

Fig. S1

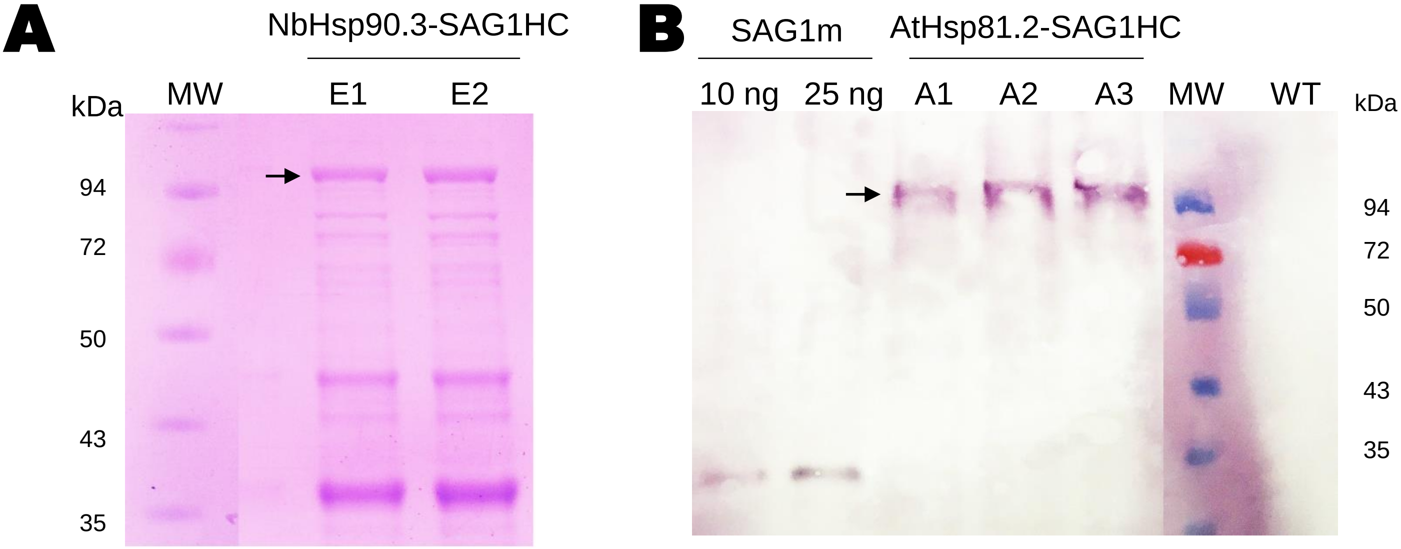


Coomassie blue

Western blot – anti-rSAG1

### Supplemental Figure 2

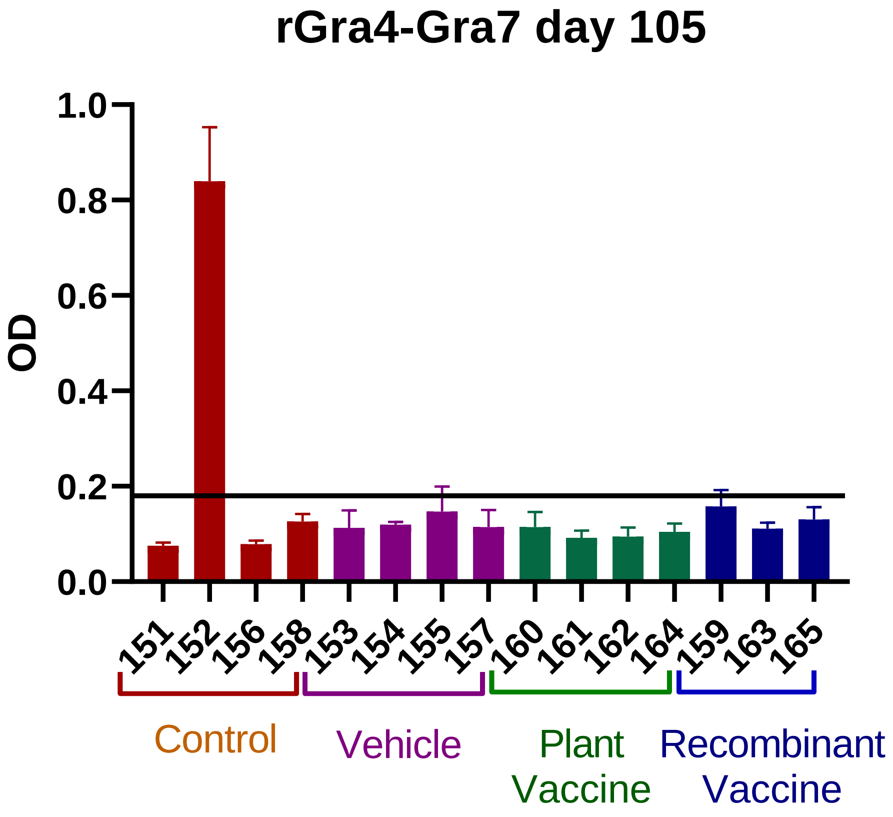

### Supplemental Figure 3

Fig. S3

**Ovine brain**

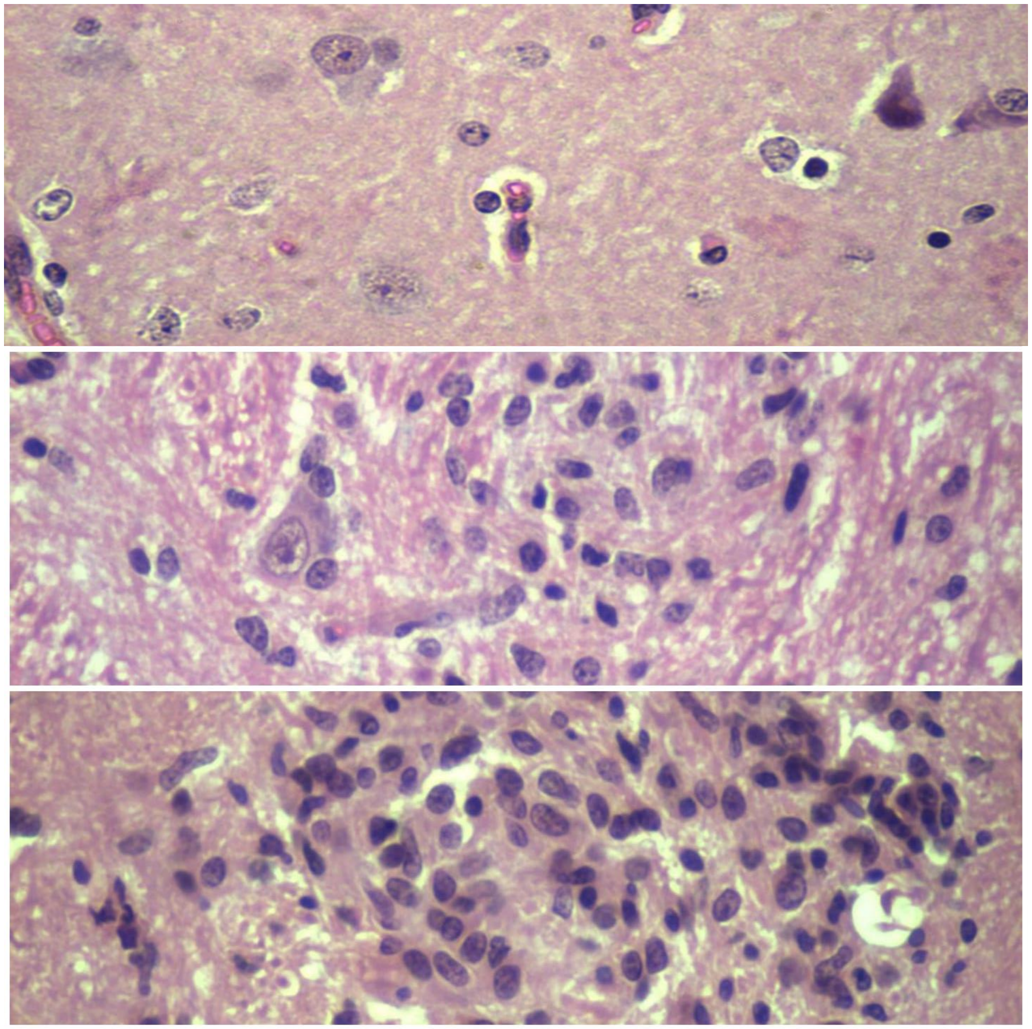

Without lessions (0)

Mild lesions (1)

Moderate lessions (2)
